# NormExpression: an R package to normalize gene expression data using evaluated methods

**DOI:** 10.1101/251140

**Authors:** Zhenfeng Wu, Weixiang Liu, Xiufeng Jin, Deshui Yu, Hua Wang, Gustavo Glusman, Max Robinson, Lin Liu, Jishou Ruan, Gao Shan

**Affiliations:** School of Mathematical Sciences, Nankai University, Tianjin 300071, P.R.China.; College of Life Sciences, Nankai University, Tianjin, Tianjin 300071, P.R.China.; School of Biomedical Engineering, Health Science Center, Shenzhen University, Shenzhen 518060, P.R.China.; Guangdong Provincial Key Laboratory of Biomedical Measurements and Ultrasound Imaging, Shenzhen 518060, P.R.China.; Institute of Statistics, Nankai University, Tianjin 300071, P.R.China.; Institute for Systems Biology, 401 Terry Avenue North, Seattle, Washington 98109, USA.; State Key Laboratory of Medicinal Chemical Biology, Nankai University, Tianjin 300071, P.R.China.

**Keywords:** gene expression, normalization, evaluation, R package, scRNA-seq

## Abstract

Data normalization is a crucial step in the gene expression analysis as it ensures the validity of its downstream analyses. Although many metrics have been designed to evaluate the current normalization methods, the different metrics yield inconsistent results. In this study, we designed a new metric named Area Under normalized CV threshold Curve (AUCVC) and applied it with another metric mSCC to evaluate 14 commonly used normalization methods, achieving consistency in our evaluation results using both bulk RNA-seq and scRNA-seq data from the same library construction protocol. This consistency has validated the underlying theory that a sucessiful normalization method simultaneously maximizes the number of uniform genes and minimizes the correlation between the expression profiles of gene pairs. This consistency can also be used to analyze the quality of gene expression data. The gene expression data, normalization methods and evaluation metrics used in this study have been included in an R package named NormExpression. NormExpression provides a framework and a fast and simple way for researchers to evaluate methods (particularly some data-driven methods or their own methods) and then select a best one for data normalization in the gene expression analysis.

## Introduction

Global gene expression analysis provides quantitative information about the population of RNA species in cells and tissues ^1^. High-throughput technologies to measure global gene expression levels started with Serial Analysis of Gene Expression (SAGE) and are widely used with microarray and RNA-seq ^2^. Recently, single-cell RNA sequencing (scRNA-seq) has been used to simultaneously measure the expression levels of genes from a single cell, providing higher resolution of cellular differences than what can be achieved by bulk RNA-seq, which can only produce an expression value for each gene by averaging its expression levels across a large population of cells ^3^. Raw gene expression data from these high-throughput technologies must be normalized to remove technical variation so that meaningful biological comparisons can be made. Data normalization is a crucial step in the gene expression analysis as it ensures the validity of its downstream analyses ^1^. Although the significance of gene expression data normalization has been demonstrated ^4^, how to select a successful normalization method is still an open problem, particularly for scRNA-seq data.

Basically, two classes of methods are available to normalize gene expression data using global normalization factors. They are the control-based normalization and the average-bulk normalization. The former class of methods assumes the total expression level summed over a pre-specified group of genes is approximately the same across all the samples. The latter class of methods assumes most genes are not Differentially Expressed (DE) across all the samples. The control-based normalization often uses RNA from a group of internal control genes (*e.g.* housekeeping genes) or external spike-in RNA (*e.g.* ERCC RNA ^5^), while the average-bulk normalization is more commonly used for their universality. Five average-bulk normalization methods designed to normalize bulk RNA-seq data are library size, median of the ratios of observed counts that is also referred to as the DESeq method ^6^, Relative Log Expression (RLE), upperquartile (UQ) and Trimmed Mean of M values (TMM) ^7^. Recently, three new methods were introduced as Total Ubiquitous (TU), Network Centrality Scaling (NCS) and Evolution Strategy (ES) with best performance among 15 tested methods ^8^. To improve scRNA-seq data normalization, Lun et al. introduced a new method using the pooled size factors (Pooled) and claimed that their method outperformed the library size method, TMM and DESeq ^9^. Bacher *et al.* addressed that using existing normalization methods on scRNA-seq data introduced artifacts that bias downstream analyses. Then, another new method SCnorm was introduced and claimed to outperform MR, Transcripts Per Million (TPM), scran, SCDE and BASiCS using both simulated and case study data ^10^.

Although many metrics have been designed to evaluate the relative success of these methods, the different metrics yield inconsistent results. The first problem in the evaluation of normalization methods is the lack of guide by a theory. In 2013, Glusman *et al.* proposed that a sucessful normalization method simultaneously maximizes the number of uniform genes and minimizes the correlation between the expression profiles of gene pairs. Based on this theory, they presented two novel and mutually independent metrics to evaluate 15 normalization methods and achieved consistent results using bulk RNA-seq data ^8^. In this study, we designed a new metric named Area Under normalized CV threshold Curve (AUCVC) and applied it with another metric mSCC (**Materials and Methods**) to evaluate 14 commonly used normalization methods using both bulk RNA-seq and scRNA-seq data from the same library construction protocol. The evaluation by AUCVC and mSCC achieved consistent results using both bulk RNA-seq data and scRNA-seq data, validating the theory proposed by Glusman *et al* ^8^. As many new normalization methods are developed, researchers need a fast and simple way to evaluate different methods, particularly some data-driven methods or their own methods rather than obtain information from published evaluation results, which could have biases or mistakes, *e.g.* misunderstanding of RLE, UQ and TMM methods ^11^. To satisfy this demand, we developed an R package NormExpression including the gene expression data, normalization methods and evaluation metrics used in this study. This tool provides a framework for researchers to select the best method for the normalization of their gene expression data based on evaluation.

## Results

### Basic concepts

In total, 14 normalization methods have been evaluated in this study. They are Housekeeping Genes (HG7), External RNA Control Consortium (ERCC), Total Read Number (TN), Total Read Count (TC), Cellular RNA (CR), Nuclear RNA (NR), the ratios of observed counts (DESeq), Relative Log Expression (RLE), upperquartile (UQ), Trimmed Mean of M values (TMM), Total Ubiquitous (TU), Network Centrality Scaling (NCS), Evolution Strategy (ES) and SCnorm (**Materials and Methods**). Currently, most methods with a few exceptions (*e.g.* SCnorm) are used to normalize a raw gene expression matrix (n samples by m genes) by multiplying a global normalization factor to each of its columns, yielding a normalized gene expression matrix (**Figure 1A**). The normalization factor in the package NormExpression is equivalent to the scaling factor in a previous study [8]. The reciprocal of normalization factor is the library size (**Figure 1B**), which represents total RNA in a library and is also named as size factor in the Bioconductor package DESeq ^6^. The definition of normalization factor in NormExpression and the definition of size factor in DESeq are different from definitions of normalization factor and size factor in the Bioconductor package edgeR ^7^. RLE, UQ and TMM in edgeR produce normalization factors to adjust library sizes, which should be used to calculate the Counts Per Million (CPM) for the normalization of gene expression data and CPM should be calculated by one million multiplying reciprocals of adjusted library sizes (**Figure 1B**). However, edgeR provides a function named calcNormFactors to produce normalization factors for library-size adjustment, which have been wrongly used for the normalization of gene expression data in many studies ^11^. The NormExpression package includes such modifications as below. Since HG7, ERCC, TN, TC, CR, NR and TU produce normalization factors by the estimation of library sizes as CPM, their normalization factors are amplified by one million for a uniform representation (**Figure 1B**). DESeq, RLE, UQ and TMM have been modified to ignore zero values for both scRNA-seq and bulk RNA-seq data and the resulting normalization factors need be further normalized by their geometric mean values (**Figure 1B**). UQ and TMM use library sizes estimated by NR. After modification, RLE is identical to DESeq. We verified that these modifications did not change the evaluation or normalization results.

**Figure 1.**
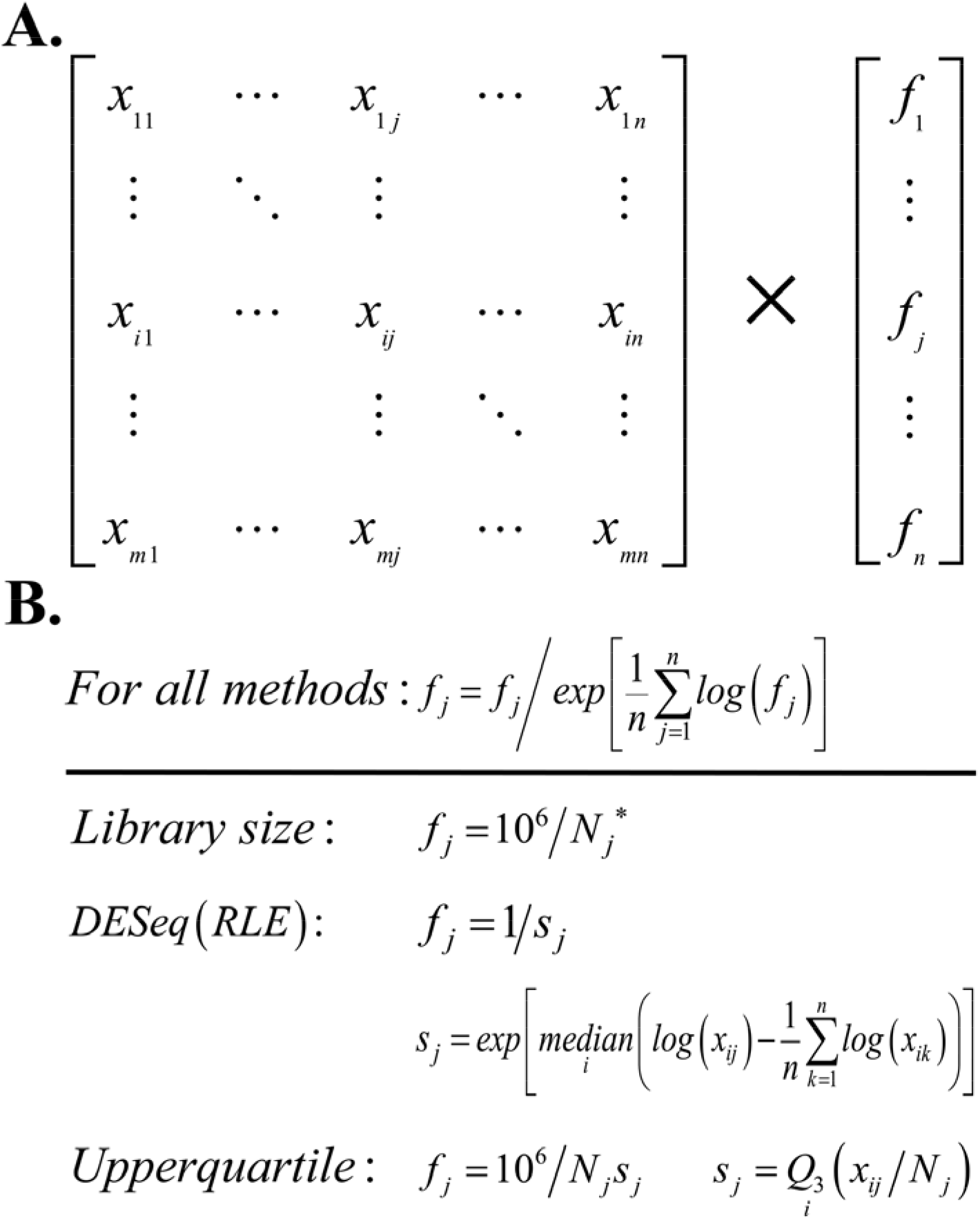
Basic concepts. (A). A raw gene expression matrix can be transformed into a normalized gene expression matrix by the multiplication of a global factor f_j_ to each column. Each column represents the expression levels of all genes from a sample and each row represents the expression levels of a gene across all samples. (B). HG7, ERCC, TN, TC, CR, NR and TU use Nj* as library sizes to calculate normalization factors. Nj represents the library size estimated by TC. DESeq, RLE, UQ and TMM have been modified in NormExpression to ignore zero values and the resulting normalization factors need be further normalized by their geometric mean values. After modification, RLE is identical to DESeq. Q3 means that about 75% of genes in the jth sample have expression levels below Q3 and about 25% have those above Q3. For all methods, log represents the natural logarithm.

### Evaluation of 14 normalization methods

In the previous study ^8^, Glusman *et al.* had quantified success of normalization methods by the number of uniform genes (**Materials and Methods**) and used the Coefficient of Variation (CV) cutoff 0.25 to determine the number of uniform genes for each method. This metric was designed based on the theory that the relative values among different normalization methods were quite stable, although the absolute number of uniform genes depended on the cutoff value. However, it is almost impossible to determine a CV cutoff for scRNA-seq data since CV in scRNA-seq data has a much larger dynamic range than in bulk RNA-seq data.

Inspired by Area Under the receiver operating characteristic Curve (AUC) ^12^, we designed a new metric named Area Under normalized CV threshold Curve (AUCVC) to evaluate 14 normalization methods using one scRNA-seq dataset scRNA663 and one bulk RNA-seq dataset bkRNA18 (**Materials and Methods**). A single housekeeping gene GAPDH was also used for comparison in the evaluation of normalization methods using bulk RNA-seq data, but it was not applicable to scRNA-seq data due to zero counts of GAPDH present in many samples. Parameter grid of non-zero ratio cutoffs (**Materials and Methods**) from 0.2 to 0.9 for scRNA663 and from 0.7 to 1 for bkRNA18 was used to produce AUCVCs of all methods (**Figure 2AB**). For each non-zero ratio cutoff, the TU method used the maximum AUCVC to determine the best presence rate (**Materials and Methods**), upper and lower cutoffs by testing all possible combinations of presence rate, lower and upper cutoffs at 5% resolution. The presence rate cutoff was tested from 0.2 to 0.6 for scRNA663 and set to 1 for bkRNA18. The lower cutoff was tested from 5% to 40% and the upper cutoff was tested from 60% to 95%. In addition, the calculation only considered each combination of lower and upper cutoffs which produced more than 1,00 ubiquitous genes (**Materials and Methods**) for scRNA663 and more than 1,000 for bkRNA18. For each non-zero ratio cutoff, NCS and ES used the presence rate, lower and upper cutoffs determined by the TU method, when it achieved the maximum AUCVC. The raw gene expression matrix (None) was also used to produce AUCVCs for comparison.

**Figure 2.**
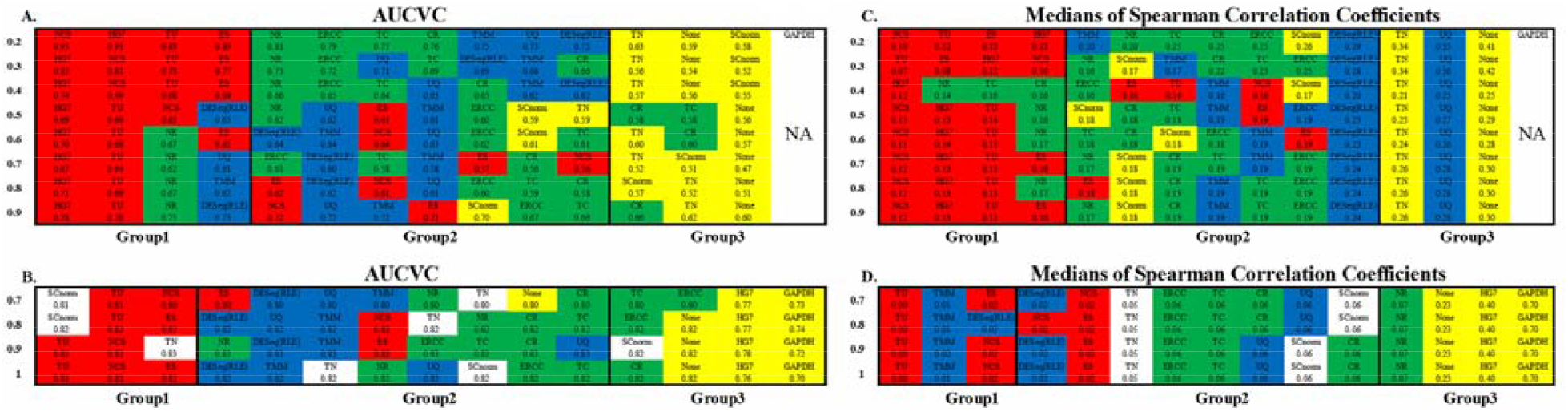
Grid of non-zero ratio to evaluate normalization methods. Grid of non-zero ratio cutoffs from 0.2 to 0.9 for scRNA663 and from 0.7 to 1 for bkRNA18 was used to produce AUCVCs All the normalization methods were classified into three groups based on their AUCVC values sorted in descending order using one scRNA-seq dataset scRNA663 (**A**) and one bulk RNA-seq dataset bkRNA18 (**B**). These methods were also classified into three groups based on their medians of SCCs sorted in ascending order using one scRNA-seq dataset scRNA663 (**C**) and one bulk RNA-seq dataset bkRNA18 (**D**). GAPDH is not applicable to scRNA-seq data due to zero counts of GAPDH present in many samples. All the numbers are accurate to two decimal places, the marginal differences are reflected by their orders.

The evaluation results using both scRNA663 and bkRNA18 showed that all methods except HG7, TN and SCnorm were classified into three groups based on their AUCVC values sorted in descending order (**Figure 2AB**). The first group including TU, NCS and ES achieved the best performances using both scRNA663 and bkRNA18. TU, NCS and ES, which had only been evaluated using bulk RNA-seq data in the previous study ^8^, were evaluated by our new metric AUCVC as the best normalization methods using both scRNA-seq and bulk RNA-seq data. The second group including ERCC, TC, CR, NR, DESeq, RLE, UQ and TMM achieved medial performances using both scRNA663 and bkRNA18. In the second group, ERCC, TC, CR and NR outperformed DESeq, RLE, UQ and TMM using scRNA663, while DESeq, RLE, UQ and TMM outperformed ERCC, TC, CR and NR using bkRNA18. The third group achieved the poorest performances, including TN, SCnorm and None for scRNA663 (**Figure 2A**) and HG7, GAPDH and None for bkRNA18 (**Figure 2B**). HG7 and GAPDH achieved the poorest performances using bkRNA18, suggesting that a predefined set of housekeeping genes may not be appropriate guides for data normalization of bulk RNA-seq data. However, it could be coincidental that HG7 was classified into the first group using scRNA663. TN outperformed the second group of methods using bkRNA18 but underperformed it using scRNA663. SCnorm was designed to improve the scRNA-seq data normalization but it performed better using scRNA-seq data than bulk RNA-seq data. Particularly, SCnorm ranked the first in the best group by its AUCVC to normalize bkRNA18 when the non-zero ratio cutoffs were set to 0.7 or 0.8 (**Figure 2B**), but ranked the last in the poorest group to normalize scRNA663 when the non-zero ratio cutoffs were set to 0.2 to 0.4 (**Figure 2A**).

The evaluation results by the medians of Spearman Correlation Coefficients between the normalized expression profiles of ubiquitous gene pairs (mSCCs) (**Materials and Methods**) were consistent with the evaluation results by AUCVC (**Figure 2**). This validated the underlying theory that a sucessiful normalization method simultaneously maximizes the number of uniform genes and minimizes the correlation between the expression profiles of gene pairs. Then, we calculated Spearman Correlation Coefficients (SCCs) bwteen all the normalization factors except that using SCnorm. Using 1- SCCs as distances, hierarchical clustering of 13 normalization factors showed equivalent classification into the same groups (**Figure 3EF**) as those (**Figure 2AB**) by AUCVC. The evaluation results by AUCVC, mSCC and hierarchical clustering can be visualized using NormExpression. From Figure 3, it can be seen that, while normalization methods except HG7, GAPDH and None perform without much differences on bulk RNA-seq data (**Figure 3BDF**), they have significant differences in the performances on scRNA-seq data (**Figure 3ACE**). This provides an explanation as to why bulk RNA-seq data and scRNA-seq data from the same samples produced different results in many biological studies. Therefore, it is more challenging to select a best method for the normalization of scRNA-seq data.

**Figure 3.**
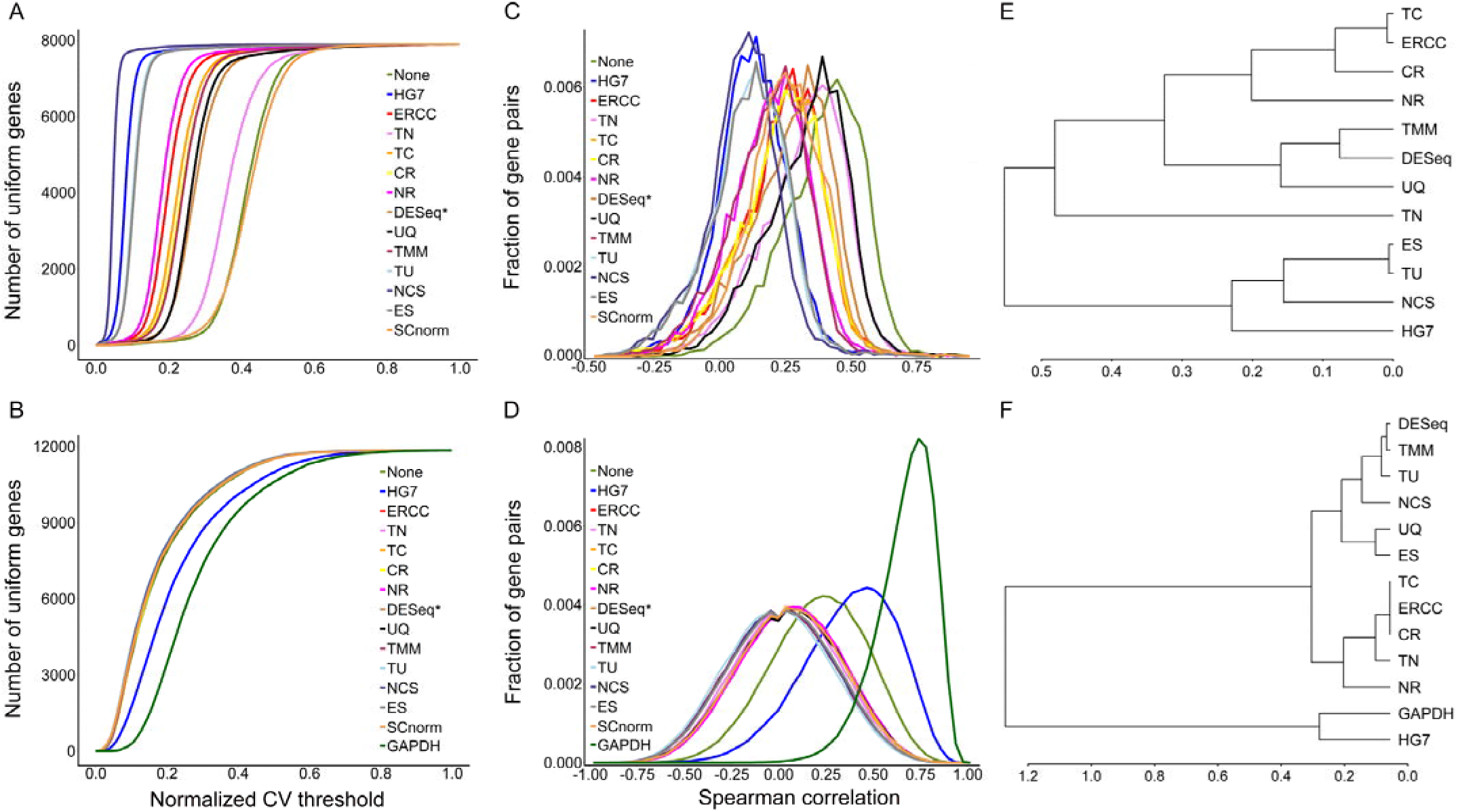
Consistency in evaluation results by different metrics. A normalization method with a higher AUCVC produced a lower median of Spearman Correlation Coefficients (SCCs) between the normalized expression profiles of ubiquitous gene pairs using both scRNA-seq (**AC**) and bulk RNA-seq data (**BD**). Using 1- SCCs as distances, hierarchical clustering of 13 methods showed equivalent classification into the same groups (**EF**) as those (**Figure 2AB**) by AUCVC. SCCs were calculated between normalization factor pairs of 13 methods. SCnorm was not applicable to be used to calculate SCCs, since it produced a factor matrix rather than a factor vector as 13 other methods.

### Implementation and availability

The gene expression data (scRNA663 and bkRNA18), normalization methods (except NCS, ES and SCnorm) and evaluation metrics (AUCVC and mSCC) have been included in the R package NormExpression. The data process in this study was provided in detail (**Supplementary file 1**). All the functions except the NCS and ES methods have been implemented in R programs for their running on R platforms of any version. The NCS and ES methods had been implemented in Perl programs with multiple dependencies on Perl modules ^8^ and were modified into a stand-alone program for Linux systems (**Supplementary file 2**).

NormExpression can be used in three ways: normalization without evaluation, normalization with simple evaluation or normalization with complete evaluation. Normalization without evaluation is suggested to use TU for users’ data, since it has been already ranked as the best method for both scRNA-seq and bulk RNA-seq data (**Supplementary file 1**). Normalization with simple evaluation by AUCVC is suggested to select the best method from 10 normalization methods using users’ data, which are HG7, ERCC (if available), TN, TC, CR, NR, DESeq (RLE), UQ and TMM. The raw gene expression matrix is also used to produce AUCVCs for comparison. TN, NCS, ES and SCnorm are not included in 10 methods, since TN and SCnorm have not shown robust performances, and NCS and ES have a similar performance of TU but are much more time consuming. Normalization with complete evaluation by AUCVC and mSCC for method selection is suggested to select the best method from at least 10 normalization methods plus TU. The normalization with simple evaluation determines the best method based on its AUCVC, while normalization with complete evaluation need consider the consistency of results using two metrics (**Materials and Methods**).

## Materials and Methods

### Datasets

In a previous study by Lin Liu *et al.* (SRA: SRP113436), 663 single-cell samples and 18 bulk samples had been sequenced using the Smart-seq2 scRNA-seq protocol. In this study, we built a scRNA-seq dataset including 653 single cells from colon tumor tissues and 10 single cells from distal tissues (>10 cm) as control. We also built a bulk RNA-seq dataset including nine samples from colon tumor tissues and nine samples from distal tissues. The cleaning and quality control of both scRNA-seq and bulk RNA-seq data were performed using the pipeline Fastq_clean [15] that was optimized to clean the raw reads from Illumina platforms. Using the software STAR ^13^ v2.5.2b, we aligned all the cleaned scRNA-seq and bulk RNA-seq reads to the human genome GRCh38/hg38 and quantified the expression levels of 57,992 annotated genes (57,955 nuclear and 37 mitochondrial). Non-polyA RNAs were not discarded to test the robustness of normalization methods, although the Smart-seq2 protocol theoretically had only captured polyA RNAs. In addition, the expression levels of 92 ERCC RNAs and the long non-coding RNA (lncRNA) MDL1 in the human mitochondrion ^14^ were also quantified. ERCC RNA had been spiked into 208 single-cell samples before library construction; the expression levels of 92 ERCC RNAs in other 455 single-cell samples and 18 bulk samples were simulated by linear regression. Finally, the two datasets were named scRNA663 (58085 × 663) and bkRNA18 (58085 × 18), and were included into the R package NormExpression.

### Normalization methods

HG7, ERCC, TN, TC, CR, NR and TU are based on a set of pre-selected genes and each of these methods uses the gene expression level summed over these pre-selected genes in a sample as the library size (**Figure 1B**) to calculate the normalization factor. The pre-selected genes used by HG7, ERCC and TU are seven housekeeping genes, 92 ERCC RNAs and ubiquitous genes (described below), respectively. Seven genes (UBC, HMBS, TBP, GAPDH, HPRT1, RPL13A and ACTB) in the HG7 method had been used to achieve the best evaluation result among those using all possible combinations of tested housekeeping genes in the previous study by Glusman *et al.* ^8^. ERCC RNA is a set of commonly used spike-in RNA consisting of 92 polyadenylated transcripts with short 3’ polyA tails but without 5’ caps ^5^. NR only counts reads which have been aligned to nuclear genomes, while CR counts reads which have been aligned to both nuclear and mitochondrial genomes. The library size estimated by TC is equal to the estimations by CR and ERCC combined. TN uses the number of all reads which can be aligned to 92 ERCC RNAs, nuclear and mitochondrial genomes.

The DESeq method was obtained from the Bioconductor package DESeq ^6^ and modified to process scRNA-seq data. RLE, UQ and TMM were obtained from the Bioconductor package edgeR ^7^ and modified to process scRNA-seq data. TU, NCS and ES were obtained from the previous study by Glusman *et al.* ^8^. Since TU sums counts of all ubiquitous genes as the library size to calculate the normalization factor, a process to identify ubiquitous genes (described below) has been integrated into the TU method: our implementation of TU maximizes AUCVC instead of the number of resulting uniform genes to determine the best presence rate, upper and lower cutoffs in the R package NormExpression. SCnorm was obtained from the Bioconductor package SCnorm ^10^.

### Uniform genes and ubiquitous genes

A gene was defined as uniform when the Coefficient of Variation (CV, **Formula 1**) of its post-normalization expression levels across all samples was not more than a cutoff. Ubiquitous genes were defined as the intersection of a trimmed sets of all samples ^8^. This trimmed set of genes were selected for each sample by 1) excluding genes with zero values, 2) sorting the non-zero genes by expression level in that sample, and 3) removing the upper and lower ends of the sample-specific expression distribution. Glusman *et al.* determined the best upper and lower cutoffs by testing all possible combinations of lower and upper cutoffs at 5% resolution to maximize the number of resulting uniform genes using one bulk RNA-seq dataset ^8^. The size of a scRNA-seq dataset is usually very large, which could result in a very small or even empty set of ubiquitous genes, since the number of ubiquitous genes depends on the sizes of datasets. To identify ubiquitous genes using scRNA-seq data, we defined a new parameter named presence rate, governing the minimal fraction of trimmed sets in which a gene must appear to be considered ubiquitous.

### Evaluation metrics

In the previous study ^8^, Glusman *et al.* designed two novel and mutually independent metrics, which were the number of uniform genes and Spearman Correlation Coefficients (SCCs) between expression profiles of gene pairs. The basic theory underlying these two evaluation metrics is that a successful normalization method simultaneously maximizes the number of uniform genes and minimizes the correlation between the expression profiles of gene pairs. In this study, we designed a new metric AUCVC instead of the number of uniform genes for method evaluation. In addtion, we randomly selected 1,000,000 ubiquitous gene pairs to calculate the mSCCs for method evaluation (**Figure 3CD**). Then, we compared the evaluation results by mSCC with those by AUCVC to assess the consistency.

AUCVC (**Figure 3AB**) is created by plotting the number of uniform genes (y-axis) at each normalized CV (**Formula** 2) cutoff (x-axis). To determine the number of uniform genes using scRNA-seq data containing a high frequency of zeros, we only considered genes with non-zero ratios not less than a cutoff. The parameter non-zero ratio of one gene should be calculated as the number of its all non-zero expression values divided by the number of total samples. Since a high or a low normalized CV cutoff produces more false or less uniform genes, it is reasonable to consider the overall performance of each method at various cutoffsettings instead of that at one specific cutoff setting. In Formula 1 and 2, symbols have the same meanings as those in Figure 1 and n* does not count zero elements in each sample.

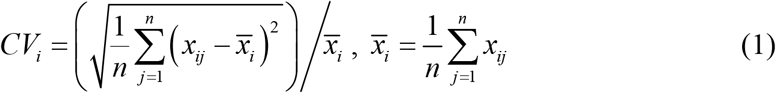

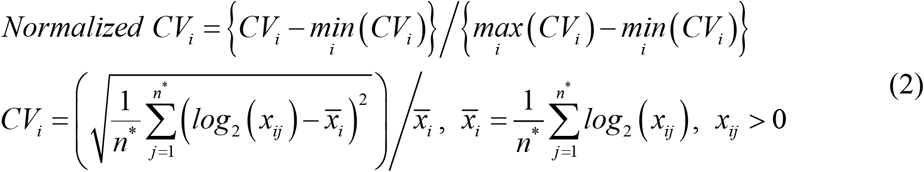

## Acknowledgments

We thank Professor Sui Huang and Dr. Joseph Zhou from Institute for Systems Biology (ISB) their hosts on Shan Gao’ visiting to ISB.

## Authors’ contributions

SG conceived this project. SG and JR supervised this project. ZW, WL and XJ performed programing. ZW and DY analyzed the data. HW prepared all the figures, tables and additional files. SG drafted the main manuscript. LL, GG and MR revised the manuscript. **All authors have read and approved the manuscript**.

